# Goliath clades and *in vivo* tracking of clonal dynamics show three phases of UV-induced skin carcinogenesis

**DOI:** 10.1101/2024.12.28.623235

**Authors:** Stanislav Avdieiev, Leticia Tordesillas, Karol Prieto, Omar Chavez Chiang, Zhihua Chen, Nihir Patel, Sofia Cordero, Luiza Silva Simoes, Y. Ann Chen, Noemi Andor, Robert A. Gatenby, Elsa R. Flores, Christopher J. Whelan, Joel S. Brown, Kenneth Y. Tsai

## Abstract

While the genetic paradigm of cancer etiology has proven powerful, it remains incomplete as evidenced by the widening spectrum of non-cancer cell-autonomous “hallmarks” of cancer. Studies have demonstrated the commonplace presence of high oncogenic mutational burdens in homeostatically-stable epithelia. Hence, the presence of driver mutations alone does not result in cancer. Here, we reveal additional forces governing the eco-evolutionary dynamics of carcinogenesis. Using a UV-driven mouse model of cutaneous squamous cell carcinoma, we tested our central hypothesis that cancer initiation occurs in three phases: 1) tissue disruption and the emergence of unusually large “goliath” clades (ecological driver), 2) clonal selection within a subset of these goliaths as evidenced by the presence of areas of unusually high local densities of cells (termed “micro-lumps”) with higher mutational burdens (evolutionary driver), and 3) emergence of macroscopic lesions. We tracked these tissue level ecological and evolutionary drivers of cancer initiation via *in-vivo* serial imaging and 3-D reconstruction of fluorescently labeled keratinocyte clades, yielding 25,085 clade measurements from 14 mice over 3 months, and 14,525 clades from 4 of these mice over 6 months. While median and mean clade sizes differed little between UV and non-UV exposure (cc. 65,000 µm^3^), our ecological survey revealed the emergence of large goliath clades (> cc. 4.2×10^6^ µm^3^), almost exclusively within UV-exposed skin. Goliath clades emerged by month 2, increased dramatically in number by months 3-4, and plateaued between months 5-6. Goliath clades arose as serpentine structures, intercalating between adjacent keratinocytes. Unexpectedly, targeted DNA sequencing revealed mutations with very low variant allele frequencies within clades, but substantial differences among clades, suggesting that positive selection for these mutations is superfluous to the development of goliaths early in carcinogenesis. scRNAseq analysis of both bulk skin and sorted clades revealed epidermal de-differentiation and immune suppression as early events. By month 7, mutational burdens were significantly larger in goliath clades, particularly for those with micro-lumps. Finally, lesions began to emerge between months 6 and 7, only in UV-exposed skin. To confirm that goliath clades are orders of magnitude more likely to spawn lesions, we randomly selected 21 goliath clades at months 6-7, prior to the time of the emergence of detectable lesions, and followed them over time. Remarkably, 2 of these developed into macroscopic lesions. Our adaptation of the Drake equation estimated the probability of this to be <10^-6^. Taken together, our results support the presence of three phases of cancer initiation, the earliest of which presage the acquisition of driver mutations and also explains why cancers are relatively rare in relation to the degree of somatic mosaicism present in UV-exposed skin.

---

Cancers have readily-defined characteristics often referred to as “hallmarks”^1^. Classically, cancer initiation is attributed to a sequential acquisition of discrete genetic events such as driver mutations. Noteworthy are the large number of mutations present in normal tissues as we age and as we are exposed to mutagens^2^. Eventually a cell achieves the right number or combination of mutations or “hits” to take it across a threshold where it can now give rise to a population of cancer cells^3^. Not fully understood is what favors the additional somatic mutations. Do they accrue from extrinsic insults to the tissue along with a regular background rate of somatic mutations, or do they accrue from intrinsic processes as some cell lineages expand at the expense of others based on the alignment of somatic mutations^4^? Both perspectives suggest that all cells of a tissue are inexorably marching their way towards cancer – some farther along than others^5^. These perspectives explain why cancer incidences increase with age, lifestyle, genetic predispositions, and inflammation and wounding. But they also suggest a concomitant breakdown in overall tissue homeostasis, composition and functioning as cells mutationally degrade^6–10^. The appearance of one cancer should portend many more.

To complement the “evolutionary perspective” of stepwise mutational accrual in cancer initiation, we suggest a parallel role for within-tissue “ecological processes” that occur through tissue turn-over, and in response to tissue damage and wounding. The need for “more to the story” comes, in part, from the empirical observation that many tissues, including skin, harbor numerous diverse somatic mutations spread across the tissue’s clones^2,11^. The vast majority of these clones never give rise to cancer. Why do they not do so? We have suggested that long runs of continuous cell division are a concomitant critical component for cancer initiation that permits the necessary driver mutations to occur within a single clone rather than spread across numerous clones. We hypothesize that it is not the average sized clade that gives rise to cancer, but rather goliath clades that emerge as part of homeostatic processes of tissue repair, regeneration and maintenance. To maintain such large clades requires long runs of cell division, just the sort that could permit the endogenous accumulation of additional somatic mutations^12^. Here we apply ecological and evolutionary principles to test whether the hallmarks of cancer are acquired in three distinct phases each with distinctive molecular and clonal properties.

To enable *in-vivo* serial assessment of clonal dynamics, we generated mice harboring the ROSA26-BRainbow2.1 cassette^13^ in combination with K14-ERT2Cre^14^ to obtain K14-CreERT2 Confetti mice. We then bred the Confetti mice with SKH1-E hairless mice (hereafter referred to as K14-Confetti mice), which are immunocompetent and susceptible to developing cutaneous squamous cell carcinoma (cuSCC) upon exposure to chronic low-dose UV irradiation^15^. Upon exposure to topical tamoxifen, one of four fluorophores is permanently expressed by K14-expressing keratinocytes and their progeny, thus comprising a group of cells expressing a single fluorophore and related by descent, hereafter referred to as clades to specifically distinguish them from genetically identical clones. Serial confocal imaging of skin on a monthly basis (Leica SP5) was followed by digital reconstruction and comprehensive morphological feature analytics. Additionally, clades were extracted at specific time points and profiled using single cell RNA sequencing and targeted panel deep DNA sequencing to reveal the molecular aspects defining the transition from normal tissue to observable cancer.

## RESULTS

### Epidermal clades form tiles with near vertical boundaries

K14-Confetti mice were tamoxifen-treated, then exposed to low-dose UVB (Oriel solar simulator; 12.5 kJ/cm^2^ weekly) intermittently for 3 months (1 hour exposure triweekly), a protocol which generates precancerous papillomas, a minority of which progress to invasive carcinomas^15^ (Fig. 1a). Fluorescently-labelled clades within the field of view were imaged by confocal microscopy (Leica SP5) and individually measured across 14 mice for 3 months (n=25,085 clade images) and, of these, 4 mice were measured for 3 additional months post-UV exposure for a total of 6 months (n=14,525 clades imaged). Five mice were sampled for scRNAseq, and 6 for deep targeted exome sequencing (Fig. 1b).

**Figure 1.**
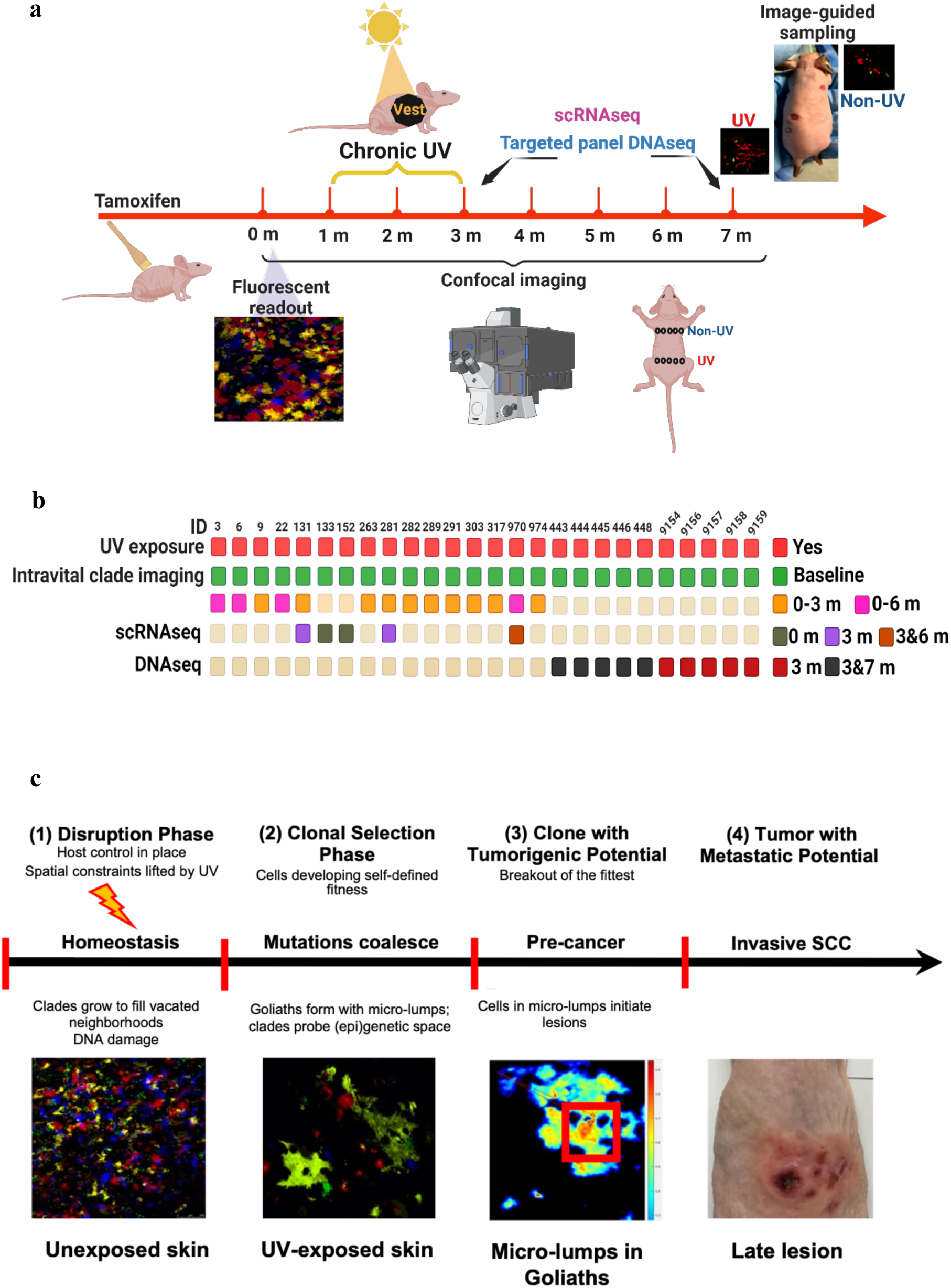
Schematic representation of the experimental approach and proposed theoretical model. **a**, Experimental setup and timeline. K14 Confetti mice were UV-irradiated for 3 months. Clones were 3-D digitized using confocal microscopy (z-stacks) and clone volumes estimated computationally. scRNAseq and targeted DNAseq were used to compare UV-exposed (EXP) vs. non-exposed (NON) epidermis vs. skin tumors. **b**, Sample heatmap of assays and metadata for individual mice used in this study. Each individual mouse ID is indicated in the top row. Mice underwent experimental procedures mentioned on the left with corresponding details on the right side of the figure. 26 mice were enrolled in this study and were exposed to UV for 3 months with the upper back shaded as an internal control. All mice were imaged at baseline (before UV exposure, month 0): 14 mice were followed by monthly imaging up to month 3 with 4 out of 14 up to month 6. Imaging data from 14 mice were used for ANOVA to investigate the interactions between UV treatment, time, and clade size. 5 mice were sampled for scRNAseq at month 0, 3 and 6, as well as for tumors at month 4, 5, and 7. 5 mice were sampled at months 3 and 7 for targeted panel DNAseq with additional 5 mice sampled at month 3. **c**, Cancer development as a process modeled in four phases. In the course of extrinsic damage, a wide variance in clone sizes begins to develop, but tissue control predominates (phase 1) until goliaths give rise to micro-lumps within which mutations accumulate, driving cell intrinsic alterations (phase 2). This, in turn results in macroscopic lesions with independent fitness functions (phase 3), further evolving into tumor with metastatic potential (phase 4).

#### Determining cut-off for clade sizes

When mice are homozygous for the Brainbow2.1 cassette, some clades express two fluorophores allowing for simultaneous visualization of the nucleus and cytoplasm when one of the two fluorophores is hrGFPII. This made it possible to delineate and count individual cells within clades (Fig. 2a). For the clade shown in Fig. 2a, the average cell size occupies 6745 µm^3^ of space. Additionally, we measured anywhere from 31 to 205 cell sizes from 10 different clades varying in size, UV exposure and month. The average volume occupied by a cell across these clades ranged from 4,410 – 8,091 µm^3^. Averaging the mean size of each clade gives 6,018 µm^3^. Rounding up to the nearest log base 2 integer gives 13. Therefore, we used 2^13^ = 8,192 µm^3^ as our cut-off for considering clade sizes in the subsequent analyses (Supplementary Table 1).

**Figure 2.**
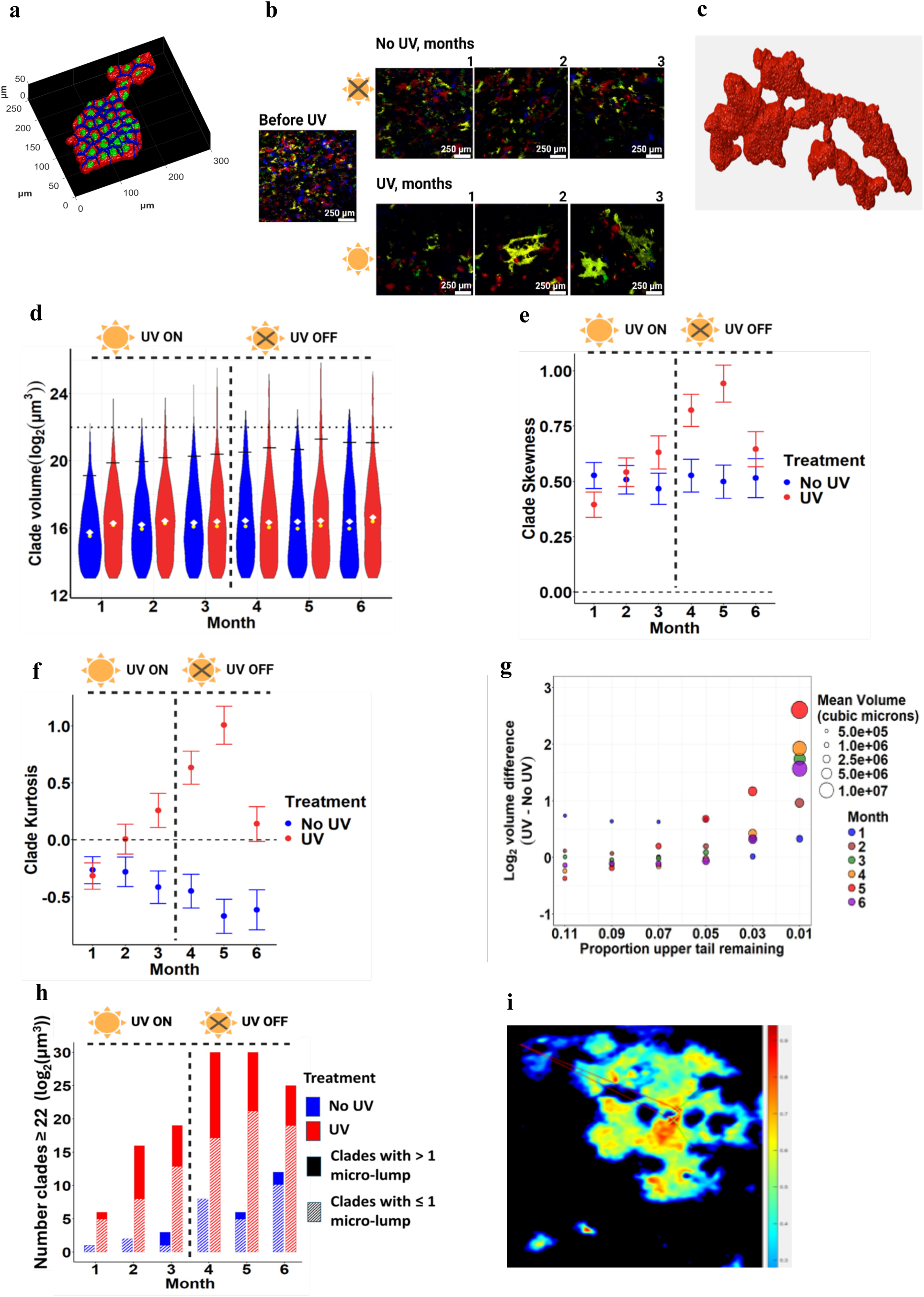
The ecology of UV effects on temporal dynamics of clade sizes and their distributions, and the emergence of goliaths. Clade morphology: **a**, single cell resolution of a clade. **b**, planar mode images of the UV-exposed and unexposed skin of the same mouse taken at month 0 (before UV) and months 1, 2, and 3 after the start of UV exposure. Note the presence of goliath clades at months 2 and 3. **c**, 3-D rendition of a clade. **d,** Distribution of clade volumes over time (n = 4 mice, months 1-6, 7,286 UV and 7,239 no-UV clades). Note that mice are exposed to UV only during months 1-3. White diamonds highlight mean clade volumes and yellow circles represent median clade volumes. tdTomato- and eYFP-expressing clades were used for the purpose of the analyses presented on the panels d-h. **e**, Both UV and no-UV clades exhibited significantly positively skewed distributions towards larger clades. UV resulted in significantly higher skew than no-UV at month 3-5 with a maximum effect at month 5. **f**, As a measure of the thickness of the tails of the distributions, no-UV always exhibited a significantly negative kurtosis (thin tails), and UV exposure resulted in a significantly positive kurtosis (wide tails) at months 3-6. Both skew and kurtosis of clades from UV-exposed skin exhibited large declines from month 5 to 6. **g**, The difference in clade volumes between those from UV exposed and clades from unexposed skin. The most pronounced effect of UV treatment occurred in the extreme right-hand tail of the clade size distribution (89-99% cutoff). The size of a bubble represents the mean clade volume at each respective time point (UV and no UV combined) (x-axis), and the y-axis gives the difference between size of UV and no-UV clades. **h**, Goliath clades accumulate over time in UV exposed skin (dashed bars) and tend to contain micro-lumps. Goliath clades with more than 1 micro-lump are depicted as solid bars. **i**, Heatmap analysis of clade cell densities demonstrate the presence of micro-lumps.

#### Clade geometries

When observed from above, clades were “pancake”-shaped and appeared as tiles in the x-y plane (looking down at the skin surface). Median sized clades (c. 68,000 µm^3^) tended to be rounded with convex edges, mean-sized clades (c. 85,500 µm^3^) began to show concavities around the edges (presumably filled in by other clades), and extremely large “goliath” clades (> 800,000 µm^3^) presented as very irregular sprawling shapes with highly irregular boundaries that in places encircled smaller clades that appeared as “holes” (Fig. 2a-c; Supplementary Movie).

#### 3D Structures of clades

In the four mice, UV-exposed clades were significantly taller than non-UV exposed clades (45.8 µm, s.e. = 0.21 µm, n=7,286, versus 40.1 µm, s.e. = 0.19 µm, n=7,239, respectively, F_1,14,513_ = 286.4, p<0.001). Clade height differed significantly contingent upon the interaction of month (1 through 6) and UV exposure (F_5,14,513_ = 14.7, p<0.001). The effect of UV was weakest at month 4 and greatest at months 1, 5 and 6, though UV exposed skin always had higher average clade heights than non-UV regardless of month (Fig. S1a; Supplementary Tables 2-3).

Because clade volume is directly related to height and area on the skin surface, we tested for verticality of clades by considering the area of each clade at its 25%, 50% and 75% height quartile from its bottom (nearest to the basal membrane) to its top (closest to the skin surface). The “walls” of clades are nearly but not quite vertical, as there is a small but significant mid-section bulge where the area of the 50% slice (8,862 µm^2^) > 75% (8,115 µm^2^) > than 25% (7,731 µm^2^; mean quartile areas of slices from n=14,525). We measured variability in the areas of different slices of a clade by creating an index of horizontal irregularity = (Area 75% - Area 25%)/Area 50%. Based on the mean areas, this index was (8115-7731)/8862, or about 4%. Generally, the mean of differences is greater than the difference of means as there can be much more variability at the individual clade level (mean of differences) than at the population level (difference of population means), and this divergence may be affected by UV. In fact, the means of the differences for UV and non-UV exposed skin were 0.052 and 0.028, respectively. The mean index of horizontal irregularity was significantly greater under UV (F_1,14521_=76.1, p<0.001). This index also increased with month irrespective of UV exposure: 0.033, 0.035, 0.040, 0.041, 0.047, for months 1, 2, 3, 4, and 5, respectively, before dropping in month 6 to 0.039; F_5,14513_=2.1, p=0.06). However, the magnitude of the difference in the index varied from month to month, leading to a significant month×treatment interaction (F_5,14513_=3.28, p<0.006). Despite the significant interaction, the index was greater for UV in each of the 6 months. Moreover, the Pearson’s correlations between the areas at the 25% and 50% slice, 25% and 75% slice and 50% and 75% slice of a clade were 0.999, 0.998 and 0.998 across clades, respectively (Fig. S1b-d; Supplementary Tables 4-6). These results suggest that, while near vertical, clades: 1) spread out more mid-height, than at their top or bottom, 2) this mid-height “bulge” increases with UV and month, and 3) despite the UV by month interaction, the effect of UV happens early in exposure and persists from month to month.

### Distribution of clade sizes is altered by UV exposure over time

#### Means, medians, skew and kurtosis

When considering clade volumes greater than 2^13^ μm^3^, the distributions of log_2_-transformed clade sizes were skewed with long tails towards larger clades (Fig. 2d; Supplementary Table 7). For the 4 mice combined over the 6 months, median (15.90 and 16.22, respectively) and mean (16.24 and 16.53, respectively) clade sizes were greater for UV-exposed than non-exposed skin: n = 7,437 versus 7,295, respectively (Fig. 2e). The taller heights of clades under UV exposure may explain much of these slight volume differences between UV and non-UV exposed skin. From these results, a range of 2^16^ = 65,536 µm^3^ to 2^16.5^ = 92,681 µm^3^ are the approximate ranges for the median and mean for clade volumes across all months and treatments. This corresponds to approximately 8-16 cells or 3-4 cell divisions from a progenitor cell.

We discretized clade volumes by rounding to the nearest integer value of log_2_ volume and subtracting 13 from this integer. All values will be ≥ 0. They represent the number of cell divisions required to generate that number of cells. Combining all clades across the 4 mice for each of the 6 months allowed us to calculate the monthly medians (always = 3, except for UV at 6 months = 4), means, skew (Fig. 2e), and kurtosis (Fig. 2f) for UV versus non-UV exposed skin. For non-UV, the means ranged from 3.18 – 3.43 for months 2 to 6, with a mean of 2.755 for month 1. For UV exposed skin, the range in means was tight (3.29 – 3.59 for months 2-6) (Supplementary Table 8).

Skew and, in particular, kurtosis highlight the effect of UV on the distribution of clade volumes (Supplementary Tables 9-12). We compared these discretized volumes to their respective Poisson distributions based on mean numbers of cell divisions (by month and UV for 12 separate comparisons). For all months, regardless of UV exposure, the observed distribution differed significantly from the expected. For all cases, the observed values at 0 and 1 cell divisions exceeded the expected, and the expected values exceeded the observed for 2, 3, and 4 cell divisions. The skew and kurtosis of a Poisson distribution is 1/λ^0.5^ and 1/λ, respectively, where λ is the mean of the distribution; therefore, expected values of skew and kurtosis are c. 0.56 and c. 0.3, respectively. For non-UV, skew across the six months was always positive, remained unchanged with month at c. 0.5, in line with a Poisson. In contrast, skew for UV changed significantly with exposure by month, rising rapidly from months 1 to 5 (reaching 0.94 at month 5) before declining significantly back towards no-UV levels at month 6 (month 5 and 6 comparison, t_1882_ = 2.62, p<0.01) (Fig. 2e). Only for months 4 and 5 did UV have a significantly higher skew than no-UV (t_2221_ = 2.84, p<0.005 and t_1909_ = 3.94, p<0.0001, respectively).

Kurtosis for no-UV was negative across all months, contrary to a Poisson distribution. In contrast, for UV, kurtosis aligns with non-UV at month 1 before rising steeply until month 5, then dropping significantly at month 6 (month 5 and 6 comparison, t_1882_ = 3.84, p<0.0002) (Fig. 2f). UV exposed skin exhibited significantly positive kurtosis for months 4 and 5 (t_1132_ = 4.38, p<0.0001 and t_854_ = 6.03, p<0.0001, respectively).

We entertain three interpretations from these distributions. First, UV has minimal effects on median and mean clade sizes. Second, the positive kurtosis and fat tail of the UV-induced distribution may result from maintaining the integrity of damaged skin. Third, the negative kurtosis for non-UV exposed skin suggests tissue level regulation to constrain the occurrence of large clades, thus shrinking the upper tail of the distribution.

#### The tail of the distributions

The above observations directed our analyses to the upper tail of the clade volume distributions. The most striking effect of UV exposure occurred in the upper tail of the distributions and in the frequencies of extremely large clades (Fig. 2g). We first considered the log_2_ difference in the mean sizes of UV vs. non-UV exposed clades at ever smaller upper portions of their respective distributions (e.g. the top 11%, 9%, 7%, 5%, 3% and 1%). From the top 5% - 11% of the distribution, the difference was roughly 0, ranging between −0.5 and 1 cell division, depending on the month and distribution cut-off. At the upper 3% the difference was always positive, though sometimes quite small, and the most extreme positive values for this difference were observed at the upper 1% cutoff. While the difference changed little at months 1 and 2, the difference increased significantly to c. 1.5 to 2 cell divisions at 1% for months 3, 4, and 6. The steepest increase reached c. 2.5 cell divisions at 1% for month 5, showing that the mean clade volume of the upper tail beyond the cutoff was over 4 times larger for UV than non-UV exposed clades. The effect of UV on the tail of the distribution is most evident in the later months (4-6 months) and at the extreme percentiles. For instance, in months 4, 5, and 6 at the upper 1%, the cutoffs for non-UV were 21.8, 21.8 and 22.3 and for UV 23.8, 24.4 and 23.9, respectively (Supplementary Table 13).

#### The first incomplete statistical moment

The first incomplete statistical moment (FISM) has been proposed as a way to differentiate between random variation (“neutrality”) in clades sizes (linear relationship between ln(FISM) and clade size) versus active selection for competitive asymmetries among clades (non-linear ln(FISM) and clade size)^16^. Competitive asymmetries are viewed as contributing towards cancer initiation. As no cancers ever emerged from the non-UV exposed skin, we might expect a difference in the shape of the FISM with clade size with UV exposure. This was not the case (Fig. S2d). The relationship between FISM or ln(FISM) was negative, significantly non-linear, and bowed outwards (getting steeper with clade size). The only notable difference occurred in the tails of the distribution, where, as noted before, the UV-exposed distribution of clade sizes extends to higher values than non-UV. In the region of extremely large clades, the relationship was very steep and linear for both UV and non-UV exposure (Fig S2d). While the overall relationship between the FISM and clade size was inconclusive in separating cancer risk between UV and non-UV exposed skin, it focused attention on the tails of the UV and non-UV clade distributions (Supplementary Table 14).

#### Goliath clades

Far in the tail of the distribution were rare, but extremely large clades. We chose log_2_ ≥ 22 as our cut-off for goliath clades which were about 64 times larger than the median clade size and c. 512 cells. This cut-off comprised approximately 1% of clades for non-UV exposed skin in all months, and UV exposed skin for months 1 and 2. For UV exposed skin at months 4 and 6 it represented about 2.5% of clades. At month 5, 3.5% of clades were goliaths for UV exposed skin. For the 4 mice, the number of goliaths was significantly higher under UV than non-UV exposure (F_1,5_=64.93, p<0.001) (Fig. 2h), and the number of goliaths varied significantly with month where months 4-6 exhibited the highest number of goliaths (F_5,5_=9.04, p<0.02). For the 4 mice over the 6 months a total of 32 and 126 goliath clades were recorded for non-UV and UV exposed skin.

Micro-lumps represented another feature associated with UV goliaths defined as localized areas with unusually high cell density as measured by a fluorescence intensity ≥ 0.9 within the clade, representing the top 10% of the signal intensity values for the total population of the yellow and red clades (Fig. 2i). Most clades exhibited no micro-lumps, and many exhibited a single micro-lump (Fig. S2e). The likelihood that a clade had one or more micro-lumps increased with clade size, with goliaths having the highest likelihood. Most strikingly, combining over months 1-6, we recorded only 4 goliath clades under non-UV with four or more micro-lumps whereas the number for UV was 42 (Fig. 2h; Supplementary Tables 15-16). Furthermore, the fraction of goliaths with four or more micro-lumps was significantly higher for UV than non-UV (0.33 versus 0.125, F_1,10_=8.13, p<0.02).

### The “Drake Equation” of cancer initiation: Do goliaths disproportionately develop lesions?

To determine whether goliath clades had a higher likelihood of producing a lesion than a randomly-selected clade, we drew an analogy with the Drake equation for estimating the probability of extraterrestrial life across the Milky Way^17^. If clades, irrespective of size, have an equal probability of spawning a cancerous lesion, what is that probability compared to the likelihood of a goliath clade spawning a lesion? For this, we required three pieces of information: 1) average number of lesions that occur on a mouse identifiably linked to a yellow or red clade, 2) average number of yellow plus red clades in the UV-exposed skin of a mouse, and 3) number of lesions assignable to a tracked cohort of yellow or red goliath clades.

For 6 mice subjected to the standard UV protocol but separate from the main experiments, we observed a total of 48 lesions (average of 8 per mouse with a range of 4 to 12). All lesions occurred in UV-exposed skin. Of these 48 lesions, 4 emerged from yellow or red clades. We rounded this up to 1 yellow or red lesion per mouse, making our calculations more conservative. Using the full experimental cohort of 14 mice at 3 months, we measured a total of 3011 yellow and red clades across the 70 sample quadrats (5 per mouse) resulting in an average of 215 yellow and red clades sampled per mouse (3,011/14). Each quadrat was 0.155 cm on a side for an area of 0.024 cm^2^ per quadrat. The total area sampled per mouse was thus 0.120 cm^2^ (5 x 0.024 cm^2^; Supplementary Table 17). This yielded an estimated 1,792 yellow and red clades per cm^2^ of UV exposed skin. The average area of UV skin for the 6 mice for which we measured the occurrence of yellow and red lesions was 9.04 cm^2^ (range of 7.91 to 9.67 cm^2^). This yielded an estimate of 16,200 UV-exposed yellow and red clades per mouse (1792 cm^-2^ x 9.04 cm^2^). With one yellow or red lesion per mouse the probability of a randomly chosen clade producing a lesion was *p* = 1/16,200 = 6.17284*10^-05^.

For the 6 mice we randomly selected 21 red or yellow goliath clades prior to the formation of lesions (3 – 4 per mouse). These were measured weekly or biweekly. There was no collective tendency for goliaths to get larger or smaller over the 6-14 weeks of measurements; goliaths shrank, grew, or remained roughly unchanged (Fig. S3; Supplementary Tables 18-19). Most notably, 2 of these 21 goliaths spawned lesions (Fig. 3). Given *p*, the probability of seeing 2 or more lesions from 21 randomly-selected clades can be determined by subtracting the probability of seeing 0 or 1 clade from a binomial expansion {1 – [(1-*p*)^21^ + 21(1-*p*)^20^*p*] = 7.996*10^-07^}. Thus, the probability of observing 2 clades of 21 becoming lesions is vanishingly small at 8.00 x10^-7^. The key to observing a clade give rise to a lesion was picking clades based on the size property of being ≥ 2^22^ µm^3^.

**Figure 3.**
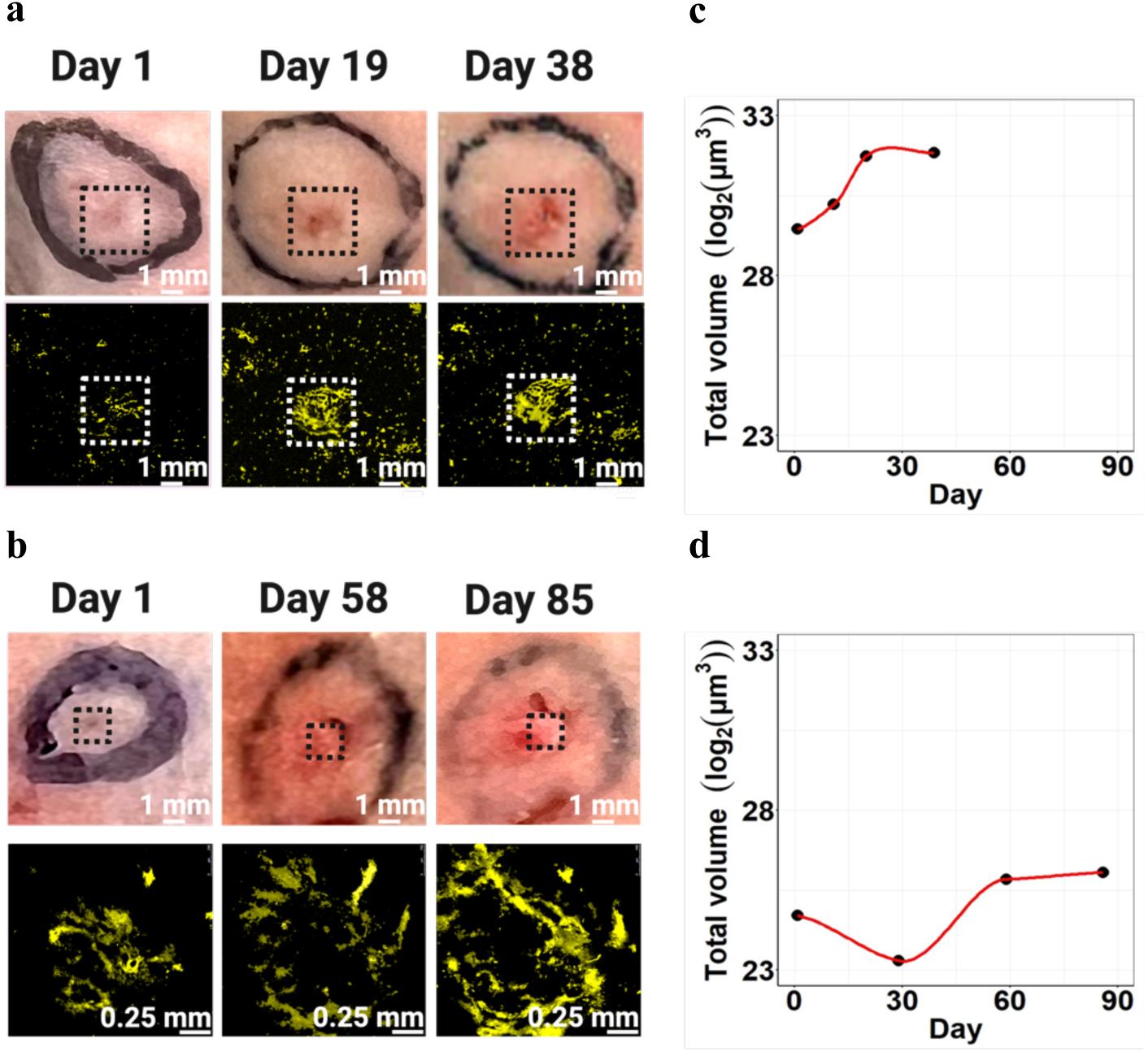
Goliath clades producing lesions representing the earliest steps of skin carcinogenesis. **a**, **b**, Goliath clades were randomly selected and marked with a tattoo line to enable tracking of the same spot (a: mouse ID 22 goliath #3; b: mouse 18 goliath #1). The clades were imaged on weekly or bi-weekly schedule. Images of the skin (top row) and corresponding 3D projections of the fluorescent images (bottom row) are shown for the same spot. We were able to observe 2 goliath-to-tumor transitions out of 21 goliath clades tracked over time. The dotted squares represent the area of developing lesion. **c, d,** The total volumes of the corresponding goliath clades were calculated based upon 3-D reconstructions and plotted as a function of time.

It may not seem surprising that larger clades of normal skin have a higher likelihood of giving rise to lesions since, but simply having more cells cannot alone explain why 2 of 21 goliaths spawned lesions. If we allow the average goliath tracked to have a size of 2^25^ --- bigger than was the case – then these 21 clades were 2^8^=256 times greater than the average clade size. If the probability of a clade producing a lesion is based on a per cell basis, then *p*’ = 0.0158 (*p*’ denoting probability scaled for size). Running this through the binomial expansion gives the likelihood of seeing 2 more or more lesions from these goliaths, based on size, of less than approximately 0.043; still unlikely. In conclusion, goliath clades had a much higher probability of producing lesions than any randomly-selected clade and this could only be partially attributed to an increased number of cells. Thus other properties of goliath clades facilitate cancer initiation.

### Mutational and transcriptional profiles of goliath clades

#### Effects of UV and clade size on mutational burdens

To identify key molecular changes associated with the development of goliaths, we performed targeted exome sequencing of 74 genes known to be involved in the development of SCC in humans and in mouse models (Supplemental Table 20) using FACS sorted cells (Fig. S4a). The Oncoplot (Fig. 4a; Supplementary Table 21) shows mutated genes in rank order of variant allele frequencies. Among the most commonly mutated genes are epigenetic regulators such as *Kmt2* family members. Mutations in well-established tumor suppressor genes such as *Trp53* are more rarely observed, in less than 30% of samples overall. *Notch* and *Fat* family members are mutated at somewhat lower frequencies in these clades and have been found in UV-exposed human skin as likely early drivers of cuSCC human skin^2^. *Prex2* mutations, which increase PI3K signaling^18^, were present in high frequency (Fig. 4a).

**Figure 4.**
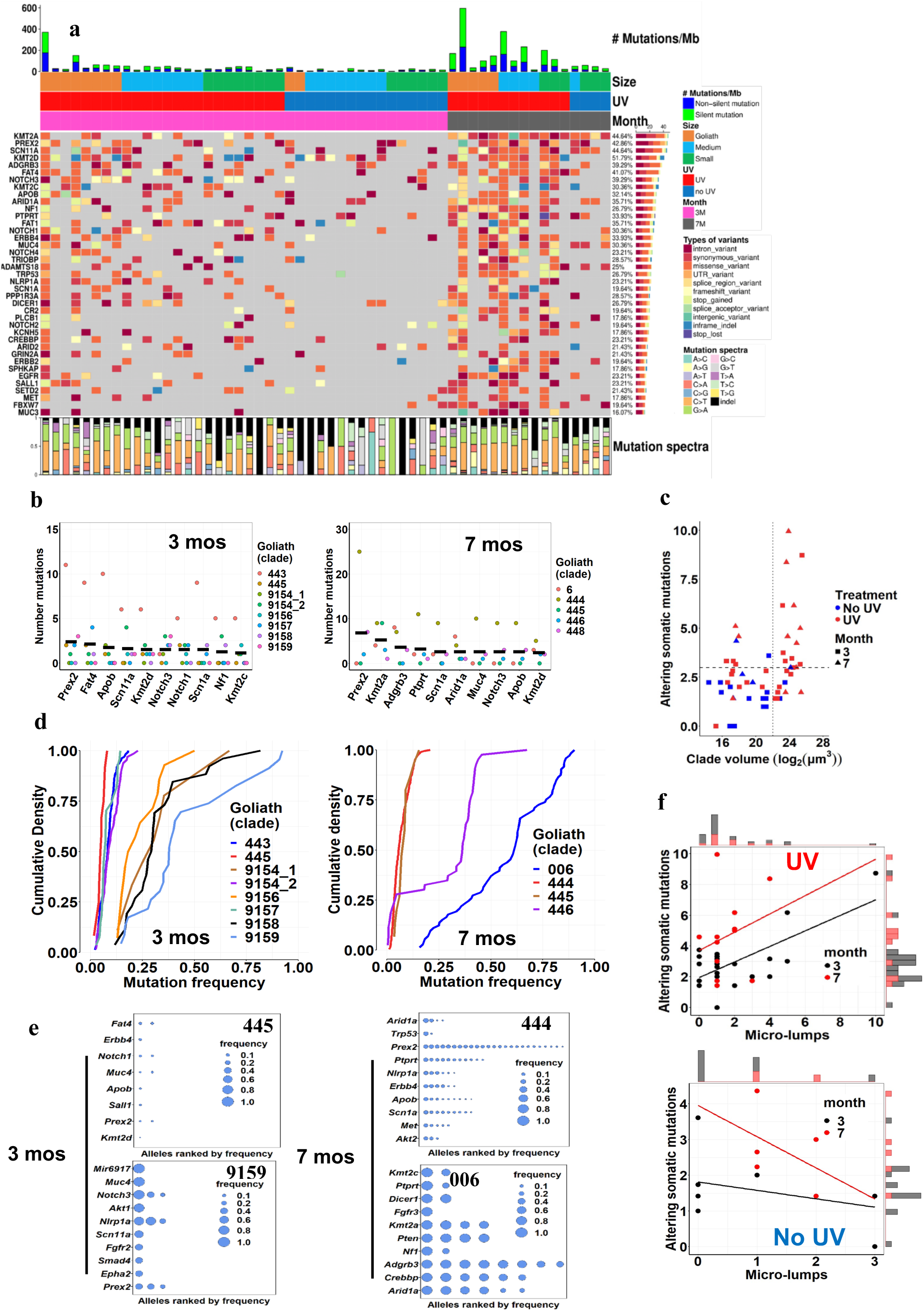
Goliath clades have diverse genetic background and do not typically possess common cuSCC drivers. a, Oncoplot summary of the mutational profile in the clades. Monochromatic clades were sampled, FACS sorted, and subjected to deep 1,000-1,500X-fold sequencing of a custom panel of 74 genes (target size ∼0.4 Mb) commonly mutated in irradiated human skin and cuSCC. UV exposure vastly increases the number of mutations observed (red vs. blue). Higher mutations loads are only observed in medium (blue) and goliath (orange) sized clades. Mutational burden is progressively increasing with the time of sampling (3 months, pink *vs*. 7 months, dark grey). **b,** Top 10 mutated genes in goliath clades sampled at 3 and 7 months from the UV exposed skin. Each dot represents mutations count per individual sample and the black bar indicates the mean. **c**, Number of altering somatic mutations increases with the size of the clade (log-transformed volume), UV treatment and sampling time (3 vs 7 months). Square root transform of the altering somatic mutations count is used for the simplicity of the analysis and indicated as mutational burden. Note that goliath-sized clades sampled from UV exposed skin later relative to the start of UV have significantly higher number of protein-altering mutations. The dotted bar at x-axis represents the arbitrary number of somatic mutations that clearly separate high burden samples from the lower ones. The bar at y-axis indicated the goliath size cutoff (2^22^ µm^3^) **d**, distribution of the mutation allelic frequencies per individual goliath clade sampled at 3 and 7 months. Each line represents an individual goliath clade. Goliath clades with high allelic frequency mutations lean to the right of the goliath clades with low allelic frequency mutations. Note the bimodal pattern in distributions reflecting tha clades arrive at the goliath size regardless of having highly selective clonal mutations. **e**, Mutations ranked by allelic frequency for the goliaths representing marginal cases on the panel d (3 months: 445 and 9159; 7 months: 444 and 006). Top 10 mutated genes are plotted (except for the clade 444, which has only 8 mutated genes in total). Bubbles represent mutations and their relative sizes reflect allelic frequency. **f**, Number of altering somatic mutations (square root transform) positively correlates with the number of micro-lumps in the UV exposed clades only.

The 10-most frequently mutated genes varied both among goliath clades and between goliath clades sampled at 3 and 7 months, as well as across clade size (Fig. 4b, Fig. S4b; Supplementary Tables 22-23). Mutational burden (shown as the square root of the number of altering somatic mutations) was greater in UV-exposed than non-UV exposed skin and was greater at 7 months than at 3 month for UV-exposed skin and for larger clades (Fig. 4c; Supplementary Table 24). Using a cut-off of 3 mutations per sample, 92% of non-exposed samples fell below this threshold, and 67% of UV-exposed samples fell above. Furthermore, for UV-exposed samples with 15 or more mutations, 80% of these were from the 7 month samples (4 months after the cessation of UV irradiation).

Interestingly, the cumulative density of variant allele frequencies (VAF) varied among goliath clades at both 3 and 7 months, with some clades reaching maximum VAF <0.025, and others reaching VAF > 0.75 at both time points (Fig. 4d; Supplementary Tables 25-26). This dichotomy reflected two extremes of mutational profiles, with some goliaths having dominant clones, and others having no discernable clonal structure, at least across established cuSCC driver genes, demonstrating that strong mutational drivers are not a prerequisite for developing goliaths (Supplementary Table 27). The relative VAF of the 10-most frequently mutated genes varied among goliath clades at both 3 months and 7 months (Fig. 4e). This further illustrated the dichotomy of goliaths with mutations of very low VAF (mice 445, 444) and those with near clonal VAFs (mice 9159, 006) at both time points. The number of altering somatic mutations increased with increasing number of micro-lumps in UV-exposed skin but decreased with increasing micro-lumps in non-UV exposed skin (Fig. 4f; Supplementary Table 28).

In examining these near-clonal clades, we found that a small clade in mouse 9159 from non-exposed skin (0.0119×10^7^ µm^3^) had mutations of likely low impact with VAF exceeding 0.94 (for example for *Cdh1*). In a small clade, the number of cell divisions was estimated to be 4 and so unlikely to represent the effects of selection primarily. In contrast, a UV-exposed goliath (mouse 006; 2.9×10^7^ µm^3^ in volume) also showed high VAF mutations, including in *Kmt2c* and *Adgrb3,* consistent with a role for epigenetic drivers through alterations in DNA methyltransferase activity.

#### Effects of UV and clade size on gene expression

We then performed single-cell RNAseq (scRNAseq) on 20 samples of dissociated skin allowing us to identify key pathways driving the transition from unexposed skin (n=10) to UV-exposed skin (n=6) and then from UV-exposed skin (n=6) to cuSCC (n=4). In the early transition following UV exposure there was upregulation of mTOR signaling, MYC signaling, and oxidative phosphorylation (Fig. 5a, left). Downregulation of p53 and interferon pathways occurred early as well and persisted in the subsequent progression to tumors (Fig. 5a, right). Importantly, some pathways, for example, oxidative phosphorylation and glycolysis, changed early but did appear to be as important in the subsequent progression to tumors, consistent with this being a molecular reflection of the multi-phasic nature of cancer development. Established oncogenic pathways, such as MYC and RAS, were clearly activated in the transition to tumors (Fig. 5a; Supplementary Table 29).

**Figure 5.**
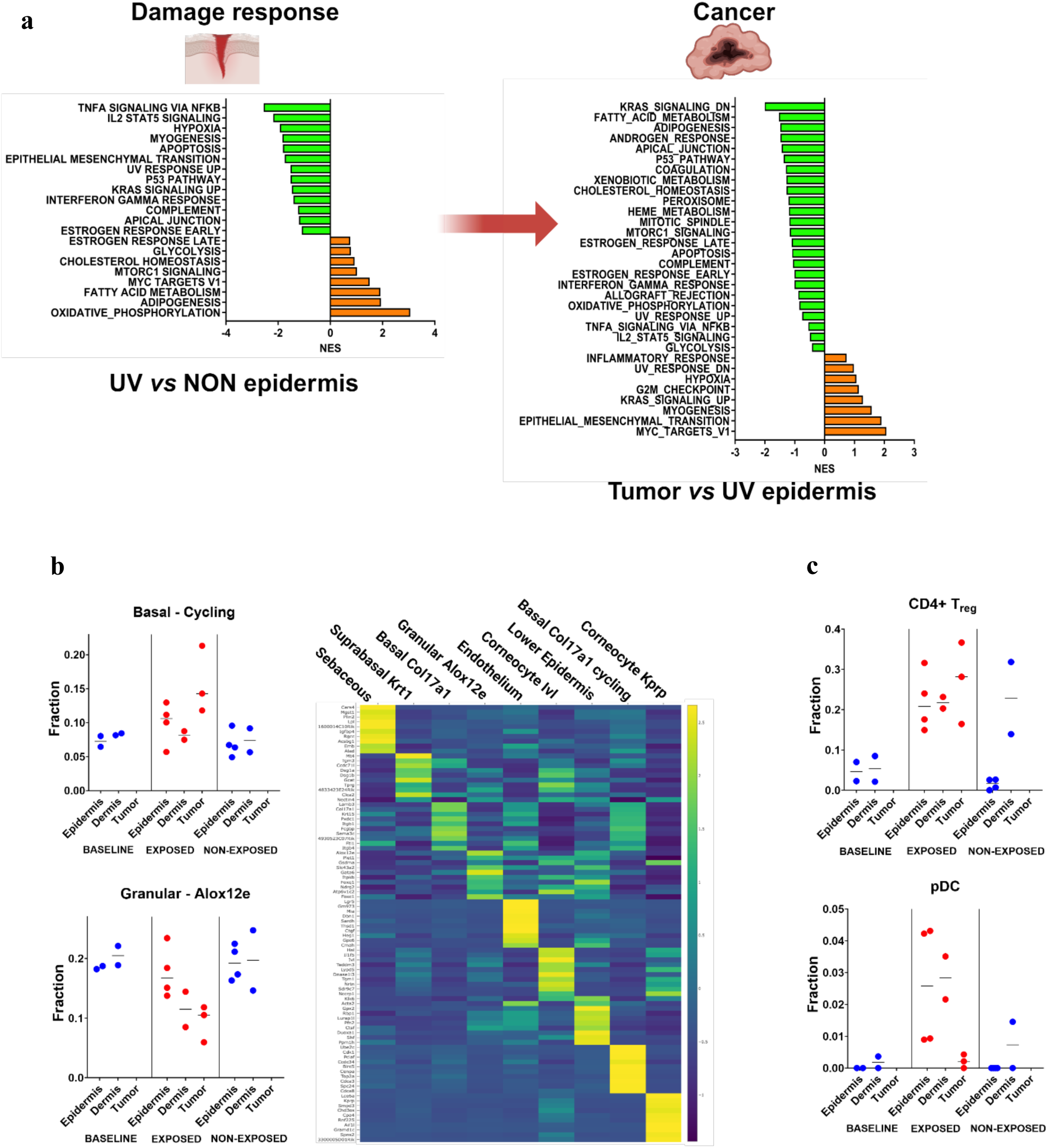
Single-cell RNAseq demonstrates dynamic transcriptional changes representing malignant transformation from normal skin to chronically exposed skin, and then to tumors. **a**, GSEA (Hallmarks) revealed up- and downregulated pathways in UV-exposed keratinocytes derived from the epidermis containing the goliaths, as well as in SCC tumors. **b**, Dynamic changes in epidermal differentiation and immune composition are plotted as proportions of total sequenced cells. Unbiased Louvain clustering revealed 9 subsets of keratinocytes defined by keratin and differentiation marker expression. **c**, The fraction of specific immune cells identified in scRNAseq data, including CD4+ regulatory T-cells (T_reg_) and plasmacytoid dendritic cells (pDC), increased following UV exposure.

#### Composition of keratinocyte and immune cell populations in normal skin, UV-exposed skin and tumors

We then tested for differences in specific keratinocyte or immune cell subsets between unexposed skin, UV exposed skin, and tumors (Fig. 5b). Subset analysis distinguished 9 major subsets of epidermal cells, including keratinocytes with varying levels of differentiation, as well as broad categories of lymphoid and myeloid (Supplementary Tables 30-32) cells.

Consistent changes were observed in UV-exposed vs. non-exposed samples, including upregulation of the fraction of cycling basal-like cells, and downregulation of Alox12e-expressing granular cells, which are found in the most superficial layers of the epidermis (Fig. 5b). We additionally observed a trend towards increased proportions of plasmacytoid dendritic cells (pDC) and of CD4+ regulatory T-cells (Tregs) in UV-exposed samples (Fig. 5c). Each of these findings highlighted key initial shifts in both keratinocyte and immune cell populations showing that UV exposure drives de-differentiation and an altered immune landscape.

## DISCUSSION

In model systems, genetically engineered mutations of established oncogenes or tumor suppressor genes alone do not confer a significant proliferative advantage in skin. For example, *Trp53*-deficient keratinocytes are outcompeted relatively quickly^6^ and *Hras*-mutant keratinocytes have no proliferative advantage with or without wounding^18^. Our interpretation of these data is that even strong driver mutations are insufficient as sole initiating events in most cancers without either powerful cooperating genetic events, chronic inflammation, or immunosuppression.

Deeply informative models such as the classical two-step carcinogenesis model clearly show that operationally, when separated, mutagenesis (e.g. *Hras* mutation) preceding inflammation gives rise to tumors^19–21^. Indeed, the stem cell of origin of tumors in this model has recently been elucidated^22^. A comprehensive CRISPR-based screen has identified that known mutational drivers of SCC, even when introduced in utero can suppress or enhance normal clonal expansion developmentally^23^. However, UV is a complete carcinogen, and exposure has profound combined inflammatory, immune, and mutagenic effects^24–26^.

Here, we probed these combined factors directly in a system that accurately captures the way in which UV exposure modulates skin ecologically and molecularly. We deployed serial imaging of lineal clades over time in the presence and absence of low-dose chronic UV exposure in an established model of cuSCC, enabling single-cell resolution 3D reconstructions of over 25,085 clades over 6 months, *through the time of macroscopic lesion formation*. Our analyses show that UV first drives an imbalance of clade size distribution with large goliath clades populating the extreme tail. In this first phase, we posit that tissue disruption from UV exposure provides a permissive environment in which **extrinsically driven** mechanisms allow for some clades to have unusually long runs of cell division and turnover. To maintain homeostatic tissue integrity under stress, cooperative dynamics between epithelial clades necessitates that some clades undergo more cell divisions than others under normal conditions.

In the second phase, we expect the emergence of **intrinsic mechanisms,** in which mutations that confer a competitive advantage allow for clade selection, with expansion of some clades at the expense of others. Even after UV-exposure is stopped, larger clades continue to provide long runs of cell division, permitting the accrual of increased molecular and genetic variability, prior to the emergence of lesions. These clades are now subject to selection for the presence of genomic drivers and pathogenic mutations. Our targeted exome sequencing reveals that there is no requirement for established mutational and pathogenic driver mutations to generate goliath clades. Our transcriptional analyses highlight key initial factors, namely the shift towards de-differentiation of epidermal keratinocytes and the increased numbers of CD4+ T_reg_ and pDC (Fig. 4c) coupled with the emergence of transcriptional signatures reflecting RAS and MYC activation.

Finally, in the third phase, we propose that cells from within extremely large clades that have undergone extensive cell divisions accumulate the **appropriate mutations** to escape local tissue control, acquire a distinct fitness function, and form tumors. In our model, unusually large clades that formed during phase 1 provided permissive environments in phase 2, resulting in the accumulation of driver mutations. Those driver mutations then served as incubators for an intra-cladal clone to emerge as a cancer in phase 3. However, the confluence of such cooperating events culminating in lesion formation is rare, as evidenced by our derivation of an equivalent “Drake equation for cancer”.

Importantly, our results complement the recent report of serial tracking of breast epithelial cells following simultaneous ablation of *Brca1* and *Trp53*^27^. We identify not only geometric constraints of clade expansion, but also identify the architecture by which clades expand, interdigitating between other clades. Our results establish that the kurtosis of clade size cumulative distributions, manifested as the fat upper tail, subsequently allow for the development of macroscopic lesions. By offering a window into the cladal dynamics, from UV exposure through macroscopic lesion formation, we document a distribution of clade sizes distinct from prior reports of mutationally-driven models, offering a complementary and distinct view into cancer initiation where ecologic and evolutionary processes set the stage for key driver mutations.

## Supporting information

Supplementary Tables Remaining (2-19, 21-24, 26-29)

## ACKNOWLEDGEMENTS

Funding support from the Melanoma & Skin Cancer, and Evolutionary Therapy Centers of Excellence at the H. Lee Moffitt Cancer Center & Research institute, and NIH NCI 5R01CA258089 (J.B. and K.Y.T.) are gratefully acknowledged. This work was additionally supported by the Molecular Genomics, Tissue Core, Analytical Microscopy, and Quantitative Imaging cores at the H. Lee Moffitt Cancer Center & Research Institute, an NCI designated Comprehensive Cancer Center (P30-CA076292).

## AUTHOR CONTRIBUTIONS

J.B. and K.Y.T. conceived the study. S.A., L.T., K.P., and O.C.-C. designed and performed experiments, imaging, irradiation, and managed animal husbandry. Z.C., Y.A.C., and N.A. performed bioinformatic analysis of single-cell RNA sequencing data. N.P. performed bioinformatic analysis of targeted exome sequencing data. S.C. and L.S.S. performed imaging statistical analyses. R.A.G. and E.R.F provided key conceptual and methodological insights J.B. and C.J.W. oversaw statistical analyses and graphical representation of data. J.B., C J.W., S.A., and K.Y.T. wrote, edited and finalized the manuscript.

## Competing interests

No competing interests are declared.

## MATERIALS & CORRESPONDENCE

Requests for materials and correspondence should be addressed to Joel S. Brown and Kenneth Y. Tsai.

## Materials and methods

### Mice

*K14-CreER* (^14^, JAX stock #005107) and *B6.129P2-Gt(ROSA)26Sor^tm1(CAG-Brainbow2.1)Cle^/J* (^28^ JAX stock #017492) mice were obtained from The Jackson Laboratory (Bar Harbor, ME). SKH1-Elite hairless mice (strain code 477) were purchased from Charles River Laboratories (Wilmington, MA). Mice used in this study were composed of males and females with mixed genetic background and aged to 90 days before commencing UV exposure. The exact age of the mice is specified in each experiment. The single-cell RNA sequencing and targeted panel DNA sequencing were performed on males aged of 90, 180, 270 and 300 days. The experiments were randomized, and no statistical methods were used to predetermine sample size. The investigators were not blinded to allocation during experiments. Mice colonies were maintained in a certified animal facility in accordance with USA guidelines. The experiments were approved by the local ethical committee (USF Comparative medicine, IACUC study IS00010270).

### Intravital laser confocal microscopy and volumetry

Our general approach utilizes the K14CreERT2; Brainbow2.1; Hairless compound mice. This mouse model has an inducible Cre-mediated recombination system driven by the human keratin 14 (KRT14) promoter ^14^, which is active in dividing basal layer epidermal keratinocytes. The Brainbow2.1 cassette ^29^ provides a multicolor fluorescent readout in which a given K14+ cell exposed to Cre will randomly express one of four fluorescent proteins (hrGFPII, EYFP, tdimer2(12) RFP, and mCerulean CFP). To ensure unambiguous distinction of the close spectral neighbors, a nuclear localization signal (NLS) was added to the carboxy-terminal end of hrGFPII.

Moreover, a membrane tethering palmitoylation sequence was introduced to mCerulean CFP. These unique features of the Brainbow2.1 cassette provide an opportunity to resolve single cell structures and enable precise optimization of morphological features for an accurate computational processing of the acquired imaging data.

Tamoxifen was applied topically on a section of the back skin at a concentration of 10 mg/ml for 3 consecutive days. Upon exposure to tamoxifen, epidermal K14+ basal cells will randomly express one of four fluorophores and this feature will be propagated into daughter cells. This allows visualization of the dynamics of clones as they arise from the basal layer over time and during tumor formation. The mice were exposed to 2.5-5 kJ/m2 of solar radiation (Oriel solar simulators, Newport) on Mondays, Wednesdays and Fridays for a total of 90 days, a dose that will produce tumors in almost all animals within the following 2 month period. A part of the mouse back skin was covered with UV-proof fabric to create UV exposed and unexposed areas within the same individual.

Mice were anesthetized with isoflurane, and intravital laser scanning confocal imaging (Leica SP5 Multiphoton) was performed in exposed and unexposed areas by random walking. Five random images were taken per each unexposed/exposed area before exposure and at 1, 2 and 3 months after the start of irradiation.

An argon laser (488 and 514 nm) and a helium-neon laser (543 and 633 nm) were deployed at 20% power to excite the above-mentioned fluorescent reporters. Each imaging area measured 1.55×1.55 mm and ranged approximately 60-100 um in depth. Chronically exposed areas tended to have higher depth. These imaging areas were sectioned optically to produce a set of z-stacks 3 µm apart starting from the basal layer up to a superficial layer of the skin. A resonant scanner frequency of 400 Hz at 1024×1024 resolution was selected as the optimal means for imaging quality/resolution and timing of the procedure. The same approach was applied to tumors imaging except for cases in which the distance between z-stacks was increased up to 10 µm to optimize timing of the procedure. Prior to recording z-stacks, the area was examined for the saturated pixels, and laser gain was adjusted appropriately. Overall, the imaging time per one field averaged approximately 10 min.

The images were then subjected to linear unmixing to ensure accurate contribution from each of the four fluorophores in the Brainbow2.1 cassette. These images were used for downstream computational processing. Noise was removed by convolving the images by image smoothing kernels and filters including Median, Gaussian, and Morphological filtering operations. A median filter was applied to the image stack with a kernel size of 3×3×3 to remove “salt and pepper” noise. A 3-D Gaussian smoothing kernel with a standard deviation of 1.0 was applied to the images to enhance image structures and reduce high-frequency intensity outliers in the images. After image filtering and de-noising, a set of morphological filtering operations were applied to the images to ensure accurate recognition of adjacent structures. This was achieved by applying border clearing operations and then applying morphological opening or closing on the grayscale image. The contrast of each image was adjusted such that all positive cells and structures were well pronounced in each fluorophore channel. The contrast was adjusted using the Percentage Linear Contrast Stretch method, with the percentage of stretched range set to 95% ^30^.

To obtain 3-D volumetric data of the clones, the images were grouped and stacked together to create a 3D image per each channel (blue, green, yellow, and red). Then, each 3D image was segmented into positively stained regions that correspond to the space occupied by a given clonal structure using the Otsu thresholding algorithm ^31^. To preserve the 3D nature of the segmented structures, both the thresholding and morphological operations were applied to the image in 3D space instead of treating each layer separately. Manual curation of the segmented and actual fluorescent images was introduced to optimize accuracy. This was achieved by numeric labeling of each segmented structure and superimposing it on the actual fluorescent image. Finally, the volumes of isolated objects of interest (clonal structures) were calculated by summing their pixel volumes.

To estimate the average volume occupied by a single cell, we selected 10 clades with clear membrane-bound distribution of the fluorophore. The individual cells were segmented, and the volumes calculated using the steps described above. We have averaged 739 individual volumes to produce the estimate of 6018 µm^3^, which we rounded up to the nearest log base 2 integer 13 (8192 µm^3^) to use as a threshold for considering clades as multicellular structures. The computational algorithm used for the image analysis is deposited in Zenodo and can be accessed through the link https://zenodo.org/records/14187948.

### Density analysis

Images underwent noise removal, contrast enhancement, and 3D segmentation as described above. The pixel intensity was calculated for the entire pool of tdTomato- and EYFP-expressing clades and scaled between 0 and 1. A cluster of pixels that have signal intensity ≥ 0.9 is considered as an area of peak density (micro-lump) and represents an area of high cell compaction. Finally, the centroids of peak density areas (x, y, and z-coordinates of the pixel that resides at the center of the dense region) are calculated and the distance from centroid to the closest and furthest clade’s edge were measured.

### Skin sampling and FACS sorting of the confetti keratinocytes

We deployed two strategies for skin sampling. Initially, for a subset of samples used for scRNAseq, we imaged the skin in exposed and unexposed areas using fluorescent confocal microscopy and biopsied using 4 mm circular blade (Accu-Punch, USA). Skin was kept afloat in the dispase (Roche Diagnostics, USA) solution for 1,5 hours. Epidermal sheets were peeled off the dermis and gently minced with a razor blade, followed by 15 min incubation in 0,25% trypsin solution (Gibco, USA) at 37C. Dermis was placed in the tissue dissociation mix (1 mg/mL collagenase type I (Sigma Aldrich, USA) and 0.1mg/mL DNase I (Roche Diagnostics, USA), minced and incubated for 1 hr at 37C. Tissues were further disaggregated by applying light pressure while pipetting and the resulting solution was applied onto 70 µM strainer. PBS/5% FBS solution was used to terminate enzymatic reaction. Samples were centrifuged at 5000 rpm for 8 min, resuspended in PBS containing 0.04% weight/volume BSA and submitted immediately for the downstream 10X single cell RNA sequencing. Alternatively, clades of the specific size category (goliath, medium, and small) were individually located in the UV exposed and non-exposed skin using fluorescent confocal microscopy and labeled with an ink. In this study, we were focused on the tdimer2(12) RFP expressing clades as an optimal means to perform cell sorting for genomics/transcriptomics analysis of the clade dynamics. Circular blade (Accu-Punch, 2 mm) was used to extract the inked skin patch containing the clade. Skin was disintegrated into dermis and epidermis as described above. The epidermal sheets were used to generate single cell suspension in FACS buffer containing EDTA, and flow sorted to collect only RFP-expressing keratinocytes representing a specific clade. Live RFP+ cells were sorted using BD FACSAria™ (BD Biosciences) at low pressure (10psi) using 130uM nozzle. Cell pellets were either preserved at −80C for the DNA sequencing or resuspended in PBS containing 0.04% weight/volume BSA and submitted immediately for the downstream 10X single cell RNA sequencing.

### Targeted panel design

A panel of 76 genes (see list below) was chosen to perform ultra-deep targeted sequencing.

ADAM29, ADAMTS18, AJUBA, AKT1, AKT2, APOB, ARID1A, ARID2, AURKA, BAI3, BRAF, CASP8, CCND1, CDH1, CDKN2A, CR2, CREBBP, CUL3, DICER1, EGFR, EPHA2, ERBB2, ERBB3, ERBB4, EZH2, FAT1, FAT4, FBXW7, FGFR1, FGFR2, FGFR3, FLG2, GRIN2A, GRM3, HRAS, IRF6, KCNH5, KEAP1, KMT2A, KMT2C, KMT2D, KRAS, MET, MUC17, MUC4, NF1, NFE2L2, NLRP1A, NOTCH1, NOTCH2, NOTCH3, NOTCH4, NRAS, NSD1, PCED1B, PIK3CA, PLCB1, PPP1R3A, PREX2, PTCH1, PTEN, PTPRT, RB1, RBM10, SALL1, SCN11A, SCN1A, SETD2, SMAD4, SMO, SOX2, SPHKAP, SUFU, TP53, TP63, TRIOBP These genes were previously reported in Martincorena et al., 2015 as genes involved in skin carcinogenesis. A custom bait capture was designed targeting the exonic sequences of these genes. Bedtools v2.26.0 was used to fetch the bed file for the genes from the gtf file or UCSC database. The overlapping regions were merged for the genes with multiple transcripts and 120 bp length sequences were generated to be used as the probes (synthesized by IDT using the IDT discovery panel).

### DNA Sequencing

Genomic DNA was extraction with QIAmp DNA Micro Kit (Qiagen). Extracted genomic DNA was then quantified with Qubit 2.0 DNA HS Assay (ThermoFisher, Massachusetts, USA) and quality assessed by Tapestation genomic DNA Assay (Agilent Technologies, California, USA). Library preparation without UMI was performed using SureSelect XT HS2 DNA Reagent Kit (Agilent), and for library preparation with UMI, xGen™ cfDNA & FFPE DNA Library Preparation Kit (IDT) was used per manufacturer’s recommendations. Exome capture was performed with IDT custom discovery pool (IDT). Library quality and quantity were assessed with Qubit 2.0 DNA HS Assay (ThermoFisher, Massachusetts, USA), Tapestation High Sensitivity D1000 Assay (Agilent Technologies, CA, USA), and QuantStudio ® 5 System (Applied Biosystems, USA). Illumina® 10-nt dual-indices were used. Equimolar pooling of libraries was performed based on QC values and sequenced on an Illumina® NovaSeq 6000 (Illumina, California, USA) with a read length configuration of 150 PE for 14 M PE reads (7M in each direction). Raw sequencing reads are accessible through the SRA database (BioProject accession PRJNA1054181).

### Clade structure and size statistical analyses

#### 3D Structures of Clade

Differences contingent upon month and treatment in both clade height and the index of horizontal irregularity were determined with the *Anova* function in the R package car ^32^.To obtain a type III sums of squares 2-way ANOVA (analysis of variance) we specified the type “III” argument in the fitted linear model. Comparisons of estimated marginal means were subsequently determined with the R package emmeans^33^.

Correlations between the quartile horizontal areas were calculated with the *correlation* function following a linear regression in base R.

#### Distribution of clade sizes

Differences in skew and kurtosis by month and UV were tested from using *SYSTAT 13*. Using the *Fitting Distributions* by month and by UV we tested for a fit of the distribution of discretized clade volumes (log_2_ −13) to a Poisson with the same mean value. The *Basic Statistics* command the mean, sample size and standard errors for skew and kurtosis by month and UV treatment. Planned comparisons month to month within UV or no-UV treatments, and UV to non-UV for a given month were conducted using t-tests using mean, sample sizes and standard errors.

#### First Incomplete Statistical Moment

We examined the relationship between the first incomplete statistical moment (FISM) of clade sizes and clade size using a linear regression of the natural log of the FISM [(ln(FISM)] and clade size with the *lm* function in base R.

#### Goliath clades

In *SYSTAT 13*, a Kruskal-Wallis 2-way nonparametric test without interaction term was used to test for the effect of month and UV on the counts of goliath clades. This was done by using the *General Linear Model* command on ranked data. A one-way Kruskal-Wallis nonparametric test was used to determine whether the frequency of goliaths with 4 or more micro-lumps differed between UV and no-UV, using the 6 months as replicates. This was done by using the *General Linear Model* command on ranked data.

### Mutational analysis

#### Non-UMI Samples

First, quality and adapter trimming were performed using trim galore and an initial quality check was performed using FastQC. The resulting trimmed reads were mapped to reference genome mm10 using bwa mem. Samtools was used to sort and index the resulting BAM files. Next, duplicate reads were marked using the Picard tool and analysis-ready BAM files were generated. Coverage at each targeted region was obtained using the Mosdepth tool. Finally, somatic mutations were detected using Mutect2 (GATK v4.1.7.0) in paired mode and were annotated using the SNPeff.

#### UMI Samples

Fastq files with UMI were first converted to unmmaped bam using FastqToSam utility (GATK v4.2.3.0), followed by UMI tag detection using ExtractUmisFromBam utility from f**gbio** toolset. The UMI tagged bam files were converted back to fastq with SamToFastq utility (GATK v4.2.3.0), mapped to reference genome mm10 using bwa mem. The UMI tagged from unmapped bam were added to mapped bam file using MergeBamAlignment (GATKv.4.2.3.0). Next, duplicate reads were marked using the Picard tool and analysis-ready BAM files were generated. Coverage at each targeted region was obtained using the Mosdepth tool. Finally, somatic mutations were detected using Mutect2 (GATK v4.1.7.0) in paired mode and were annotated using the SNPeff.

### OncoPlot

The putative somatic mutations that were annotated as PASS by Mutect2 were considered for further analysis. Finally, R package ComplexHeatmap (version 2.13.1) was used to generated oncoplots.

### Mutation Signature Analysis

An allele frequency threshold of 0.5% was applied to capture the final set of protein-altering mutations. For all samples, mutation signature analysis was carried out using an R package, musicatk using SBS96 motifs. Finally, the detected signatures were compared against Cosmic signature V2 and homology between two signatures was calculated using cosine similarity. The comparison plots are generated for top5 signatures for all samples as well individually for UV treated and control samples.

https://github.com/campbio/musicatk

https://cancer.sanger.ac.uk/signatures/signatures_v2/

SBS96 - Motifs are the six possible single base-pair mutation types times the four possibilities each for upstream and downstream context bases (464 = 96 motifs)

Reference to Packages Used in Analysis.

1. Trim_galore (v0.6.4): https://www.bioinformatics.babraham.ac.uk/projects/trim_galore/
2. BWA mem (v0.7.17) https://github.com/lh3/bwa
3. FastQC (v0.11.9): https://www.bioinformatics.babraham.ac.uk/projects/fastqc/
4. Samtools (v1.10.2): http://www.htslib.org/
5. GATK, Mutect2, Picard (v4.1.7.0): https://github.com/broadinstitute/gatk
6. Fgbio(v1.3.0) https://github.com/fulcrumgenomics/fgbio
7. Mosdepth (0.2.7): https://github.com/brentp/mosdepth
8. SNPeff (v4.3): https://pcingola.github.io/SnpEff/
9. OncoPrint : https://www.cbioportal.org/faq#what-are-oncoprints

*Cumulative density functions of mutation frequencies* were calculated with the *stat-ecdf* function in the R^34^ package ggplot2^35^. Differences among the cumulative density distributions were tested with the *k-samples* function in the R package dgof^36^.

### 10X single cell sequencing

Single-cell RNA-sequencing was performed using the 10X Genomics Chromium System (10X Genomics, Pleasanton, CA). The cell viability and counts were obtained by AO/PI dual fluorescent staining and visualization on the Nexcelom Cellometer K2 (Nexcelom Bioscience LLC, Lawrence, MA). Cells were then loaded onto the 10X Genomics Chromium Single Cell Controller at a concentration of 1,000 cells/µl in order to encapsulate 2,000 cells per sample. Single cells, reagents, and 10X Genomics gel beads were encapsulated into individual nanoliter-sized Gelbeads in Emulsion (GEMs), and reverse transcription of poly-adenylated mRNA was performed inside each droplet at 53°C. The cDNA libraries were then completed in a single bulk reaction by following the 10X Genomics Chromium NextGEM Single Cell 3’ Reagent Kit v3.1 user guide, and 50,000 sequencing reads per cell were generated on the Illumina NextSeq 2000 instrument. Prior to the data analysis, a mouse custom reference genome was built by adding *hrGFPII*, *EYFP*, *tdimer2(12) RFP*, and *mCerulean CFP* following the instruction of building a custom reference on the 10x genomics web. The Demultiplexing, barcode processing, alignment, and gene counting were performed using the 10X Genomics CellRanger v7.1.0 software, The results of the analysis were visualized using the 10X Genomics Loupe browser v6.4.1.

### Single cell RNAseq quality control and cell typing

A two-component computational tool *ISCVA*, Interactive Single Cell Visual Analytics ^37^, was used to perform scRNA-seq analyses, cell type classification, curation, and visualization. It consists of two major components. One includes the back-end scripts, functions utilizing R packages commonly utilized by single cell communities while the second component uses front-end JavaScripts to enable interactive investigation. Briefly, the R package, Seurat, was used to process the aggregated transcript count matrix generated from the 10X Cell Ranger pipeline. A total of 66199 cells from 20 samples were sequenced. One sample was excluded for low number of called cells (<1000). From the remaining 19 samples, 47226 cells passed QC (mitochondria content < 20%, number of unique genes > 500, and number of detected molecules > 1000) were used in this analysis. In our scRNA-seq analysis the 10X Chromium Single-Cell 3′ Library, Prep Kit was used for the sample preparation. Regularized negative binomial regression, along with a second linear regression against mitochondria read percentage as implemented in SCTransform, was used to normalized and adjust for cell-to-cell technical variations. A two-stage clustering process was performed. The first stage was for broad cell type identification while the second stage was for cell sub-population identification within each of the three subsets: lymphoid, myeloid, and melanoma plus fibroblast cells, respectively. The approach of the two-stage clustering and how the number clusters are selected for investigation are described in our previous work^37,38^. Briefly, at each stage, we used the unsupervised clustering algorithms, Louvain at different clustering resolutions (with resolution parameters set at 0.6, 0.8, 1, 1.2, 2, and 4) and Infomap, in the principal component analysis (PCA) space to identify cell groups among all cells that passed QC. Followed by clusters generated at different resolutions as described above, we curated cell types against multiple reference panels provided in SingleR and other known signatures ^37,39^ and visualized them interactively through ISCVA. In addition, the R package, SingleR, was used to generate cell type candidates for each cell using multiple reference panels. The finest level of clusters, which we could align to currently known cell types, would be the number of clusters to be selected as the curated cell types in ISVCA. For some of the cell types, such as tumor cells, there would be multiple clusters due to biological heterogeneity. Each cell group (or main category) is then assigned a broad cell category based on the combination of SingleR predictions – using majority vote using the cell type in mousernaseq panel and followed by curation. In ISCVA, multiple resolutions of these clusters are available for visualization and future investigation. The second stage of clustering and curation mapped out the substructure of the myeloid and lymphoid cell populations. The myeloid cell analysis included all cells that fell under the categories of pDC and macrophage/monocyte/DC, while the lymphoid analysis included all T and NK cells. The second stage analyses generated unsupervised clustering of the sub-populations of cells and then annotated these clusters according to their distinguishing gene expression, using published markers on myeloid and lymphoid cell subpopulations as a guide. The raw and processed data files can be accessed through GEO database (accession number GSE276788).

### Statistical methods

Means and medians of the clade volumes, as well statistical significance of the clade volume differences between various time points and treatments were calculated using Systat v.13^40^.

Cumulative density functions of mutation frequencies were calculated with the *stat-ecdf* function in the R^34^ package ggplot2^35^. Differences among the cumulative density distributions were tested with the *k-samples* function in the R package dgof^36^.

Differences contingent upon month and treatment in both clade height and the index of horizontal irregularity were determined with the *Anova* function in the R package car^32^. To obtain a type III sums of squares 2-way ANOVA (analysis of variance) we specified the type “III” argument in the fitted linear model. Comparisons of estimated marginal means were subsequently determined with the R package emmeans^33^.

We examined the relationship between the first incomplete statistical moment (FISM) of clade sizes and clade size using a linear regression of the natural log of the FISM [(ln(FISM)] and clade size with the *lm* function in base R. Correlations between the quartile horizontal areas were calculated with the *correlation* function following a linear regression in base R.

## Notes

### Competing Interest Statement

The authors have declared no competing interest.

