## Supplementary Data and Figures for "Goliath clades and *in vivo* tracking of clonal dynamics show three phases of UV-induced skin carcinogenesis"

[1]Cancer Biology and Evolution Program
[2]Department of Integrated Mathematical Oncology
[3]Department of Tumor Microenvironment & Metastasis
[4]Department of Biostatistics and Bioinformatics
[5]Admera Health, 126 Corporate Blvd, South Plainfield, NJ 07080.
[6]Department of Metabolism and Cancer Physiology
[7]Department of Pathology
[8]Melanoma & Skin Cancer Center of Excellence

H. Lee Moffitt Cancer Center and Research Institute, Tampa, FL, 33612 USA

**SUPPLEMENTARY TABLE LIST**

| | |
|---|---|
| Supplementary Table 1 | Data for single cell volume calculation |
| Supplementary Table 2 | Clade heights over time (4 mice, month 1-6) |
| Supplementary Table 3 | Clade heights over time (14 mice, month 1-3) |
| Supplementary Table 4 | Pearson's correlations using the areas at 25% and 75% slice from the bottom of the clade |
| Supplementary Table 5 | Pearson's correlations using the areas at 25% and 50% slice from the bottom of the clade |
| Supplementary Table 6 | Pearson's correlations using the areas at 50% and 75% slice from the bottom of the clade |
| Supplementary Table 7 | Distribution of clade volumes over time |
| Supplementary Table 8 | Difference in clade volumes between UV and non-UV skin |
| Supplementary Table 9 | Clade skewness over time (4 mice, months 1-6) |
| Supplementary Table 10 | Clade kurtosis over time (4 mice, months 1-6) |
| Supplementary Table 11 | Clade skewness over time (14 mice, months 1-3) |
| Supplementary Table 12 | Clade kurtosis over time (14 mice, months 1-3) |
| Supplementary Table 13 | Clade fractions as a function of the volume (4 mice, month 1-6) |
| Supplementary Table 14 | First incomplete statistical moment as a function of clade volume (4 mice, months 1-6) |
| Supplementary Table 15 | The distribution of clade micro-lumps in UV vs non-UV skin (4 mice, months 1-6) |
| Supplementary Table 16 | Number of goliaths with micro-lumps over time |
| Supplementary Table 17 | Average number of lesions; Average number of red/yellow lesions; back area estimate for the Drake equation |
| Supplementary Table 18 | Volume dynamics over time of the goliaths that produced lesions |
| Supplementary Table 19 | Volume dynamics of all the goliath clades tracked over time |
| Supplementary Table 20 | Targeted DNAseq gene panel and sequencing probes |
| Supplementary Table 21 | Oncoplot summary of the mutational profile in the clades |
| Supplementary Table 22 | Top 10 mutated genes in goliath clades from UV skin |

| | |
|---|---|
| Supplementary Table 23 | Top 10 mutated genes in the small and medium clades from UV vs non-UV skin |
| Supplementary Table 24 | Number of altering somatic mutations and clade size |
| Supplementary Table 25 | Clades sorted by VAF; Cell number and division estimate |
| Supplementary Table 26 | Distribution of the mutation allelic frequencies in the goliath clades |
| Supplementary Table 27 | Mutations ranked by allelic frequency for the goliaths representing extremes of VAF |
| Supplementary Table 28 | Relationship of altering somatic mutations and micro-lumps |
| Supplementary Table 29 | GSEA (Hallmarks) comparing clades from UV vs non-UV skin, and tumors vs. clades from UV skin |
| Supplementary Table 30 | scRNAseq epithelial fractions and heatmap table |
| Supplementary Table 31 | scRNAseq lymphoid fractions |
| Supplementary Table 32 | scRNAseq myeloid fractions |

### SUPPLEMENTARY FIGURES

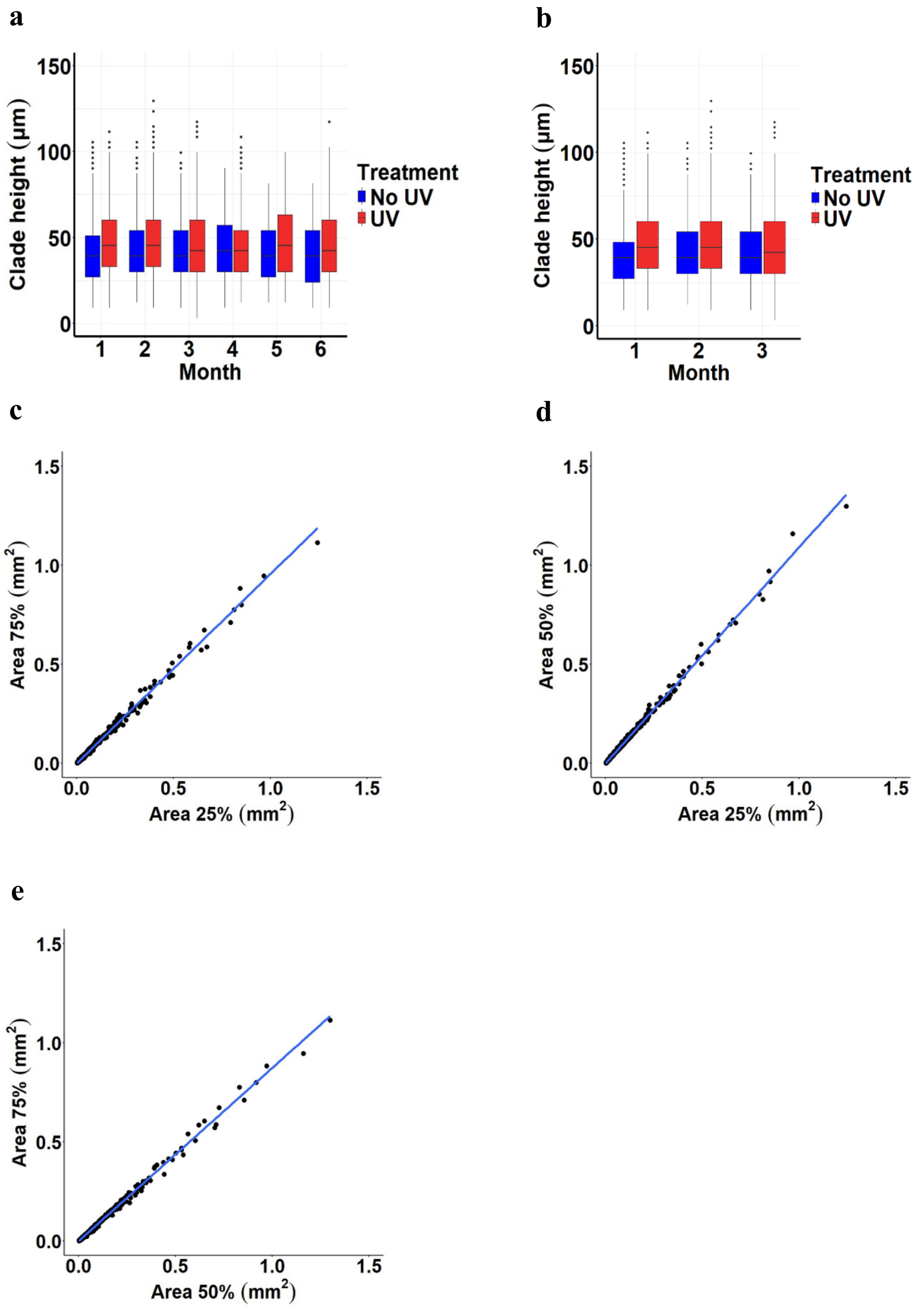

**Figure S1. Clade morphology.**

**a**, Effect of UV on clade height was analyzed using a cohort of 4 mice (month 1-6, 7,286 UV and 7,239 no-UV clades). **b**, Effect of UV on clade height was analyzed using a cohort of 14 mice (month 1-3, 11,808 UV clades and 13,277 no-UV clades) The clades under UV are taller than non-exposed regardless of the month ($p<0.001$). **c, d, e,** The areas of the clades at optical section representing its 25%, 50% and 75% height quartile from the bottom were calculated and the Pearson's correlations analysis performed using the areas at 25% and 75% slice, 25% and 50% slice, and 50% and 75% slice (Pearson's correlation 0.998, 0.997 and 0.996, respectively). The data represent 4 mice at month 1-6 with 14525 clades total. The clades are near vertical with some overlap mid-height.

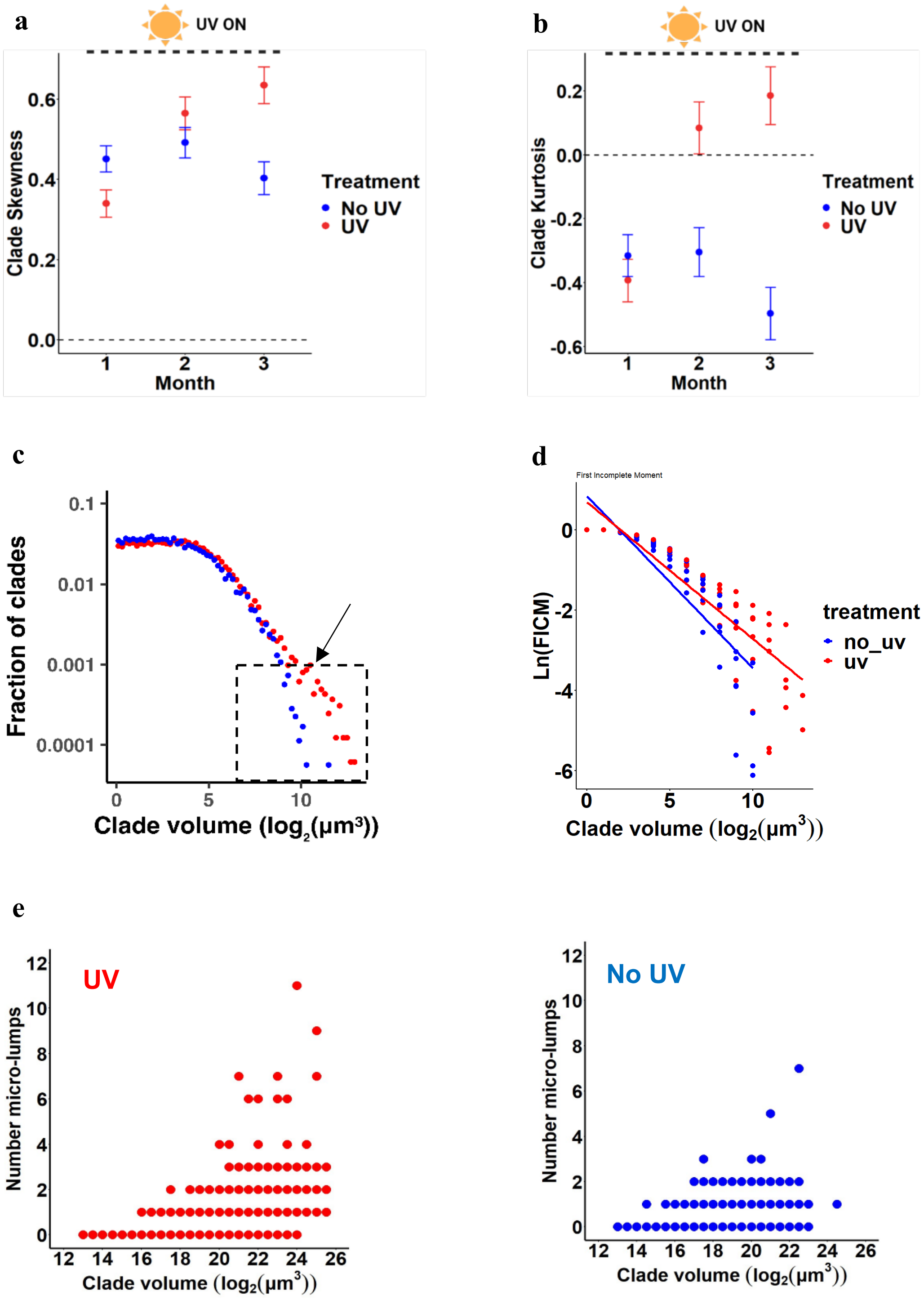
a
UV ON
Clade Skewness
0.6
0.4
0.2
0.0
1
2
3
Month
Treatment
No UV
UV
b
UV ON
Clade Kurtosis
0.2
0.0
-0.2
-0.4
-0.6
Month
Treatment
No UV
UV
c
Fraction of clades
0.1
0.01
0.001
0.0001
0
5
10
Clade volume (log2(μm³))
d
First Incomplete Moment
Ln(FICM)
treatment
no_uv
uv
Clade volume (log2(μm³))
e
UV
Number micro-lumps
Clade volume (log2(μm³))
No UV
Number micro-lumps
Clade volume (log2(μm³))

**Figure S2. The ecology of UV effects on temporal dynamics of clade sizes, their distributions, and micro-lumps development within goliath clades.**

**a**, Effect of UV treatment on clade sizes were tested on the cohort of 14 mice at month 1, 2, and 3 (11,808 UV clades and 13,277 no-UV clades). Both UV and no-UV exhibit significantly positively skewed distributions towards larger clades. UV results in significantly higher skew than no-UV at month 3. **b,** Using the same dataset presented in **a**, clade kurtosis was calculated. Clades from no-UV exposed skin always exhibit a significantly negative kurtosis (thin tail) while clades from UV exposed skin results in a significantly positive kurtosis (fat tail) at months 2 and 3. **c**, Fraction of the clades as a function of volume (log-transformed) (n = 4 mice, month 1-6, 7,286 UV and 7,239 no-UV clades). Note the divergence of the tails for UV (red) and no-UV (blue) clades. **d**, First incomplete statistical moment as a function of clade volume (n = 4 mice, month 1-6, 7,286 UV and 7,239 no-UV clades). The relationship is highly non-linear for both UV (coefficient of squared clade volume term = -0.026, t = -5.47, P = 5.68e-07) and no-UV clades (coefficient of squared clade volume term = -0.073, t = -9.3, P = 2.63e-13); **e,** The distribution of micro-lumps in no-UV *vs*. UV-exposed skin. Uniform pixel intensity settings were applied to 7,286 UV and 7,239 no-UV clades. Density analysis was performed to identify areas of the peak intensity *a.k.a.* micro-lumps (≥0.9 on the scale 0-1). Micro-lumps tend to cluster within extreme goliath clades in the UV exposed skin.

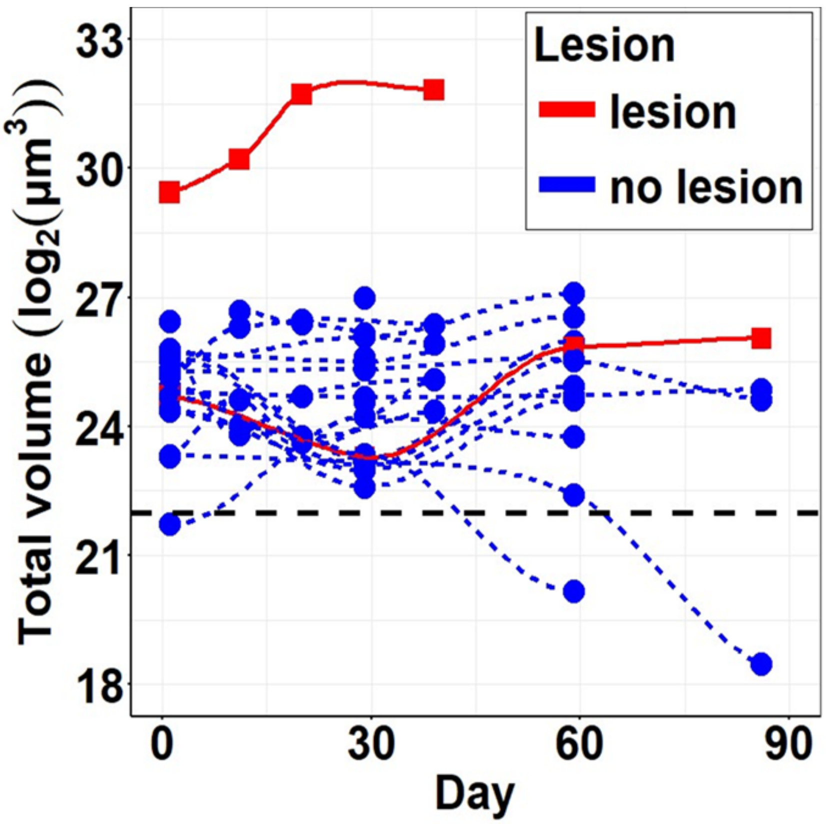

Lesion
lesion
no lesion
Total volume (log2(μm3))
33
30
27
24
21
18
0
30
60
90
Day

**Figure S3. Goliath clades have various volume dynamic over time.**

21 goliath clades have been haphazardly selected and marked with a tattoo line to enable tracking of the same spot. The clades were imaged on weekly and/or bi-weekly schedule. The total volume of the corresponding goliath clades were estimated (Methods) and plotted as a function of time. Two goliaths that produced lesions are marked in red.

a

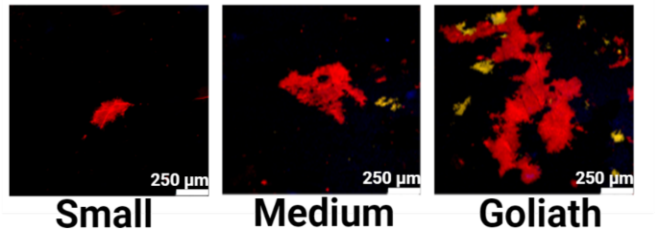
250 µm
250 µm
250 µm
Small
Medium
Goliath

b

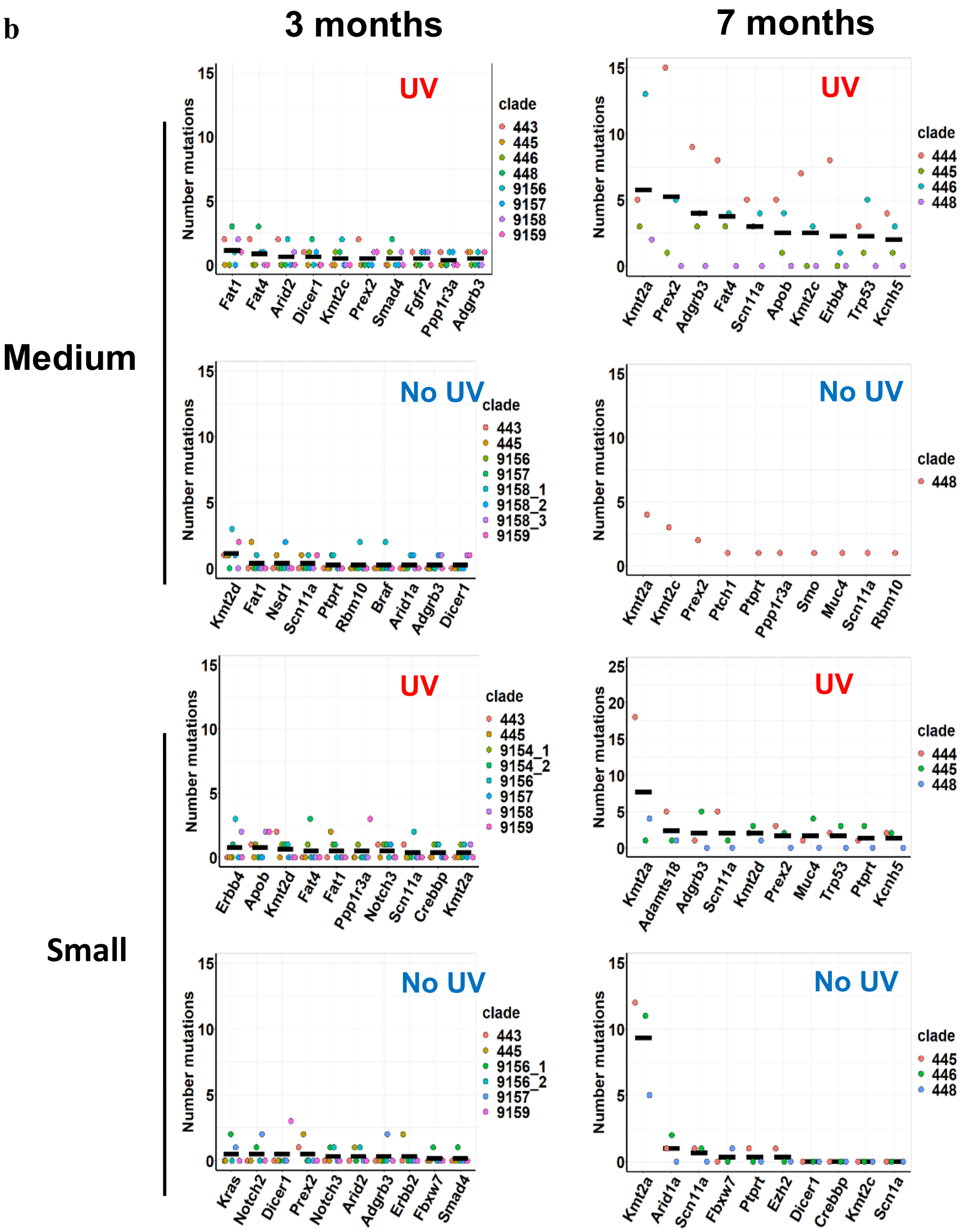
3 months
7 months
Medium
Small
UV
No UV
Number mutations
clade
443
445
446
448
9156
9157
9158
9159
444
9158_1
9158_2
9158_3
9154_1
9154_2
9156_1
9156_2
Fat1
Fat4
Arid2
Dicer1
Kmt2c
Prex2
Smad4
Fgfr2
Ppp1r3a
Adgrb3
Kmt2a
Scn11a
Apob
Erbb4
Trp53
Kcnh5
Kmt2d
Nsd1
Ptprt
Rbm10
Braf
Arid1a
Ptch1
Smo
Muc4
Notch3
Crebbp
Adamts18
Kras
Notch2
Erbb2
Fbxw7
Ezh2
Scn1a

**Figure S4. Representative images of the sampled clades and the most mutated genes per individual clade.**

**a,** representative image of the clades of three different size categories that were extracted from the epidermis at 3 and 7 months post chronic UV exposure, FACS sorted, and sequenced using targeted gene panel. **b,** Top 10 mutated genes are demonstrated for the monochromatic clades clustered according to the size category and UV treatment. Each dot represents mutational counts per individual clade and the black bar indicates the mean.
